# Assaying how the success of phagocytosis depends on the mechanics of a large viscoelastic target

**DOI:** 10.1101/545004

**Authors:** M. Davis-Fields, L. Bakhtiari, K. Kovach, V. D. Gordon

## Abstract

The state of the art does not provide a method for determining how the success of phagocytosis depends on the mechanics of a target that is much larger than the phagocytosing cell. We have developed such a method. We vary the elastic moduli of millimeter-sized abiotic gels that contain fluorescent beads to act as tracers for phagocytosis. We isolate human neutrophils, expose them to gels for one hour, and then measure what percentage of neutrophils contain beads – this is our metric for successful phagocytosis. Both increased polymer concentration in agarose gels and increased crosslinking density in alginate gels are associated with decreased success of phagocytosis. When we plot the percentage of neutrophils containing beads as a function of the gel elastic modulus, we find that data from both alginate and agarose gels collapse onto the same curve. This demonstrates the utility of our method as a way of measuring how the viscoelastic mechanics of a large target impact the success of phagocytosis.

## Statement of Significance

Bacterial biofilms are viscoelastic materials made of bacteria embedded in a matrix of extracellular polymers and proteins. Biofilm infections resist clearance by the immune system and have been shown to consist of multiple discrete aggregates, each approximately 100 μm in diameter, that are densely surrounded, but not entered, by neutrophils that are approximately 10 μm in diameter. Neutrophils are phagocytic immune cells that readily engulf bacterial cells that are not embedded within a biofilm community. For any phagocytic cell and any much-larger target, whether pieces of the target can be detached for subsequent engulfment must depend on both the force exerted by the cell and on the mechanics of the target. Our measurements also show that the success of phagocytosis depends strongly on elastic modulus for the range of elastic moduli that we previously measured for biofilms re-grown from clinical isolates (~0.05 – 10 kPa).

## Introduction

Neutrophils are short-lived phagocytic cells that are an integral part of the innate immune system and the most abundant white blood cell in humans ^2–4^. During the initial inflammatory response to infection, neutrophil migrate from the bloodstream into infected or damaged tissue where they can clear away pathogens and debris ^5^. Phagocytosis is one of a neutrophil’s main ways of clearing pathogens ^6^. Others have shown that neutrophils, which are approximately 10μm in diameter, are unable to engulf rigid polystyrene particles, with a modulus of about 3,000 MPa, that are over about 10μm in diameter ^7^. In addition, size as well as other geometric characteristics are also known to influence phagocytosis by macrophages, which are another type of phagocytic immune cell ^8–11^. Thus, the size of the target is a geometric, and therefore a physical, limitation on phagocytosis.

Single bacteria are approximately 1μm in diameter and are readily engulfed by neutrophils. However, biofilm infections in soft tissue are made up of aggregates that are typically 100μm in size – far too big to be cleared by neutrophils if the biofilm acts like a solid, rigid target as described above ^7, 12–14^. Indeed, biofilm infections are typically resistant to clearance by neutrophils ^15–17^. Imaging studies of chronic biofilm infections show bacterial aggregates that are densely surrounded by neutrophils that do not enter the aggregates and are unable to clear the infection ^12, 13^

However, biofilms are not rigid solids but are composite viscoelastic materials. We have shown that biofilms re-grown from clinical isolates have elastic moduli in the range ~0.05-10 kPa ^18^. We also found that *in vivo* evolution in chronic Cystic Fibrosis (CF) infections changed the production of matrix polymers in a way that promoted mechanical toughness ^18^. Others have found that neutrophils can exert an attractive stress of about 1kPa during phagocytosis^7^; this is consistent with the forces exerted by macrophages and blood granulocytes during phagocytosis^8, 19–21^. Thus, the mechanical stress exerted by neutrophils and other phagocytic immune cells during phagocytosis is within the range of elasticity characterizing biofilms. Therefore, it is possible that variation in biofilm elasticity could impact biofilms’ resistance to clearance by neutrophils and that the mechanical robustness promoted by changes in polymer production in clinical isolates could increase the biofilm’s ability to avoid phagocytosis by neutrophils.

Indirect support for this idea comes from a recent finding by others. *Trichomonas vaginalis* is a pathogen too large for neutrophils to engulf whole. It was recently shown that neutrophils can detach and engulf fragments of *Trichomonas vaginalis*, using a process called trogocytosis ^*22–25*^. Neutrophils can also kill cancer cells using trogocytosis ^26^. The nibbling process of trogocytosis shows that neutrophils have the ability to remove small pieces from a large target before engulfing those pieces ^26^. Phagocytosing *Dictyostelium* and *Entamoeba* can also break off and ingest small pieces of a target ^27, 28^.

We expect that how successful neutrophils are at detaching pieces of biofilm for subsequent engulfment must depend on the viscoelastic mechanics of the biofilm material. However, prior studies examining physical limitations of phagocytosis have focused on the size and shape of rigid targets ^7–11^. One earlier study using macrophages from mice showed that the stiffness of target particles could act as a cue for phagocytosis, with stiffer particles being more likely to be engulfed – however, the particles used were 6 μm or less in diameter, and therefore well below the limiting size for phagocytosis ^29^. Protocols for determining how a large, viscoelastic target’s mechanics impacts its susceptibility to phagocytosis, by neutrophils or by any other phagocytic cell, do not exist in the current state of the art. This prevents both our basic understanding of neutrophil phagocytosis and the development of novel anti-biofilm treatments that could weaken biofilms to make them more susceptible to clearance by the immune system.

To fill this gap and therefore have the ability to elucidate the mechanical limitations on phagocytosis, we developed a method to determine the impact of viscoelastic mechanics on neutrophils’ ability to successfully engulf pieces of centimeter-sized, viscoelastic targets. For two different hydrogel chemistries, we find that as the elastic modulus of the target gel varies across the range we previously measured for biofilms, the success of engulfment changes, from about 30% success at low elasticity to essentially 0% success at high elasticity. This is the first demonstration of a mechanical limit on phagocytosis. Thus, we demonstrate that a mechanical property of the target is correlated with the degree of successful engulfment, and that this experimental design is appropriate for probing the effects of target viscoelasticity on phagocytosis by neutrophils. Therefore, this work constitutes both a step forward in basic understanding and a methodological development that can be used to advance the field.

## Results

### Measurements of Neutrophil Interactions with Biofilms

We used time-lapse phase contrast microscopy to image neutrophils interacting with biofilm-like aggregates of bacteria that were an order of magnitude bigger than the bacteria. This was intended primarily to assess whether or not neutrophils could remove individual bacteria from the aggregates, and secondarily to allow us to estimate an approximate timescale for this process. On several occasions, we indeed saw neutrophils remove and engulf one or two bacteria out of an aggregate (Supplementary Movies 1-3). Because these bacteria were initially embedded in a biofilm-like matrix, these are examples of neutrophils causing localized cohesive failure in biofilms.

To estimate the frequency characterizing the distortion imposed on the biofilm aggregates by neutrophils, variously-sized protrusions across five different biofilm-attacking neutrophils were observed and the time that elapsed during each retraction of a protrusive appendage was measured. The inverse of this time was taken to be the frequency characterizing the biofilm distortion that could be imposed by the retraction of that particular protrusion. The retraction of five different protrusions from four different neutrophils were measured. On average, this frequency was 0.06 Hz, with a standard error of the mean of 0.1 Hz; 0.06 Hz is approximately equivalent to an angular frequency of 0.38 radians/sec. The average distance covered by protrusions was 2.91 μm, and the estimated average speeds were 0.26 μm/s and 0.18 μm/s for extension and retraction respectively. Previous researchers have leading edge of a neutrophil protrusion wrapping around a bead at 0.1 μm /sec, which is a value comparable to the speeds we measure ^7^.

In the next section, we describe the mechanical properties of hydrogels that we measured using oscillatory bulk rheology. At the low frequencies characterizing neutrophil-imposed deformations, the moduli have no significant dependence on frequency for all gels used (Figure 1A; Figure 2A). Therefore, the moduli reported by low-frequency bulk rheology are the moduli that could be relevant for a neutrophil attempting to detach a piece from a larger gel.

### Mechanical Properties of Hydrogels

To assess the mechanical properties of the hydrogels to be used as targets for phagocytosis, we used oscillatory bulk rheology. To control for temperature-dependence of the elastic modulus^30^, all rheology was done after the gels were allowed to sit at 37°C in an incubator for one hour to ensure the properties being measured were the same as those in the experiments in which we exposed gels to neutrophils at 37°C. We used two hydrogel types, agarose and alginate, to be able to vary the elasticity across comparable ranges using two different chemical compositions.

#### Agarose gels

Agarose gel is a product of agar or agar-bearing marine algae. It is a linear polymer with alternating D-galactose and 3,6-anhydro-L-galactose units. It is well-known that increasing the concentration of polymer in a gel will increase the gel’s elastic modulus^31^. We varied agarose concentrations from 0.3% - 2% and the resulting range of elastic moduli, ~0.1 to ~10 kPa, roughly covered the range of elastic moduli that we measured previously for biofilms grown from clinical isolates of *Pseudomonas aeruginosa* ^18^. The value of the elastic modulus at the midpoint of the plateau region in strain sweeps is the value we will use to characterize these gels’ mechanics, unless the material was already yielding for the lowest strain used – in that case, the value of the elastic modulus that was measured at the lowest strain will be used (Figure 1B). For each concentration of agarose used, three replicate gels were made and measured. The measured values were then averaged to determine the elastic modulus for that agarose concentration (Table 1).

**Table 1.**

|  | **Agarose Gels** |  |  |  |  |
| --- | --- | --- | --- | --- | --- |
| Agarose concentration | 0.30% | 0.50% | 1.0% | 1.5% | 2.0% |
| Elastic Shear Modulus<br>(Standard Error of the<br>Mean) in Pascals | 125 (3) | 370 (60) | 2,250 (70) | 5,600 (300) | 10,500<br>(700) |

**Figure 1.**
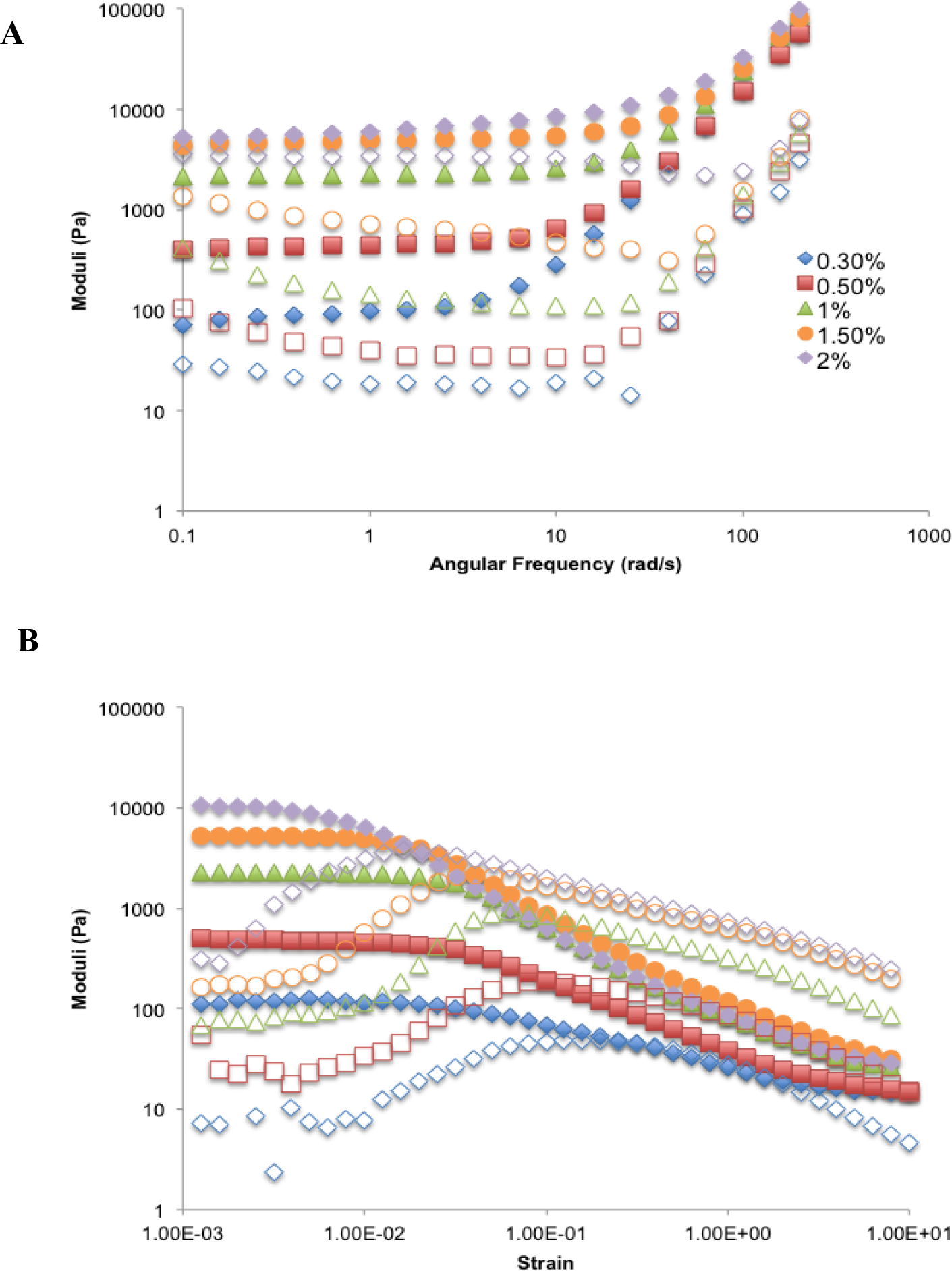
Representative frequency sweep and strain sweep curves of agarose hydrogels. (A) Frequency and (B) strain sweeps for one set of agarose gels ranging in concentration from 0.3% agarose to 2% agarose. Frequency sweeps were done at 1% strain and strain sweeps were done at 3.14 radians/s. The elastic moduli (G’) are shown with solid symbols and the viscous moduli (G”) are shown with hollow symbols of corresponding shape and color.

#### Alginate

Alginate polymers are made up of β-D-mannuronate (M) and α-L-guluronate (G) residues. Calcium ions, which are divalent, crosslink the G residues of alginate polymers. Increasing the crosslinking density, by increasing the concentration of calcium ions, increases the elastic moduli of alginate gels ^32^. This is consistent with what we found using bulk rheology (Figure 2). We used calcium concentrations of 10mM, 20mM, and 30mM which resulted in elastic moduli of the alginate gels ranging from 0.1kPa-4.5kPa. For alginate gels made with 10mM calcium, the elastic modulus measured was frequency dependent only at frequencies more than an order of magnitude greater than the ~0.06 Hz (or 0.38 radians/sec) frequency characterizing neutrophil phagocytosis. At low frequency, the elastic modulus for alginate gels made with 10 mM calcium remained under 100Pa, which is an order of magnitude lower than the elastic modulus measured for alginate gels made with 20mM calcium (Figure 2B. The reported elastic modulus for the 10mM alginate gels used for the remainder of this paper will be the value at the midpoint of the plateau region of the strain sweep. For each concentration of calcium used, three replicate alginate gels were made and measured. The measured values were then averaged to determine the elastic modulus for alginate gels made with that concentration of calcium (Table 2).

**Table 2.**

|  | **Alginate Gels** |  |  |
| --- | --- | --- | --- |
| Calcium concentration | 10 mM | 20 mM | 30 mM |
| Elastic Shear Modulus (Standard Error of the Mean) in Pascals | 13 (1) | 1500 (500) | 4500 (300) |

**Figure 2.**
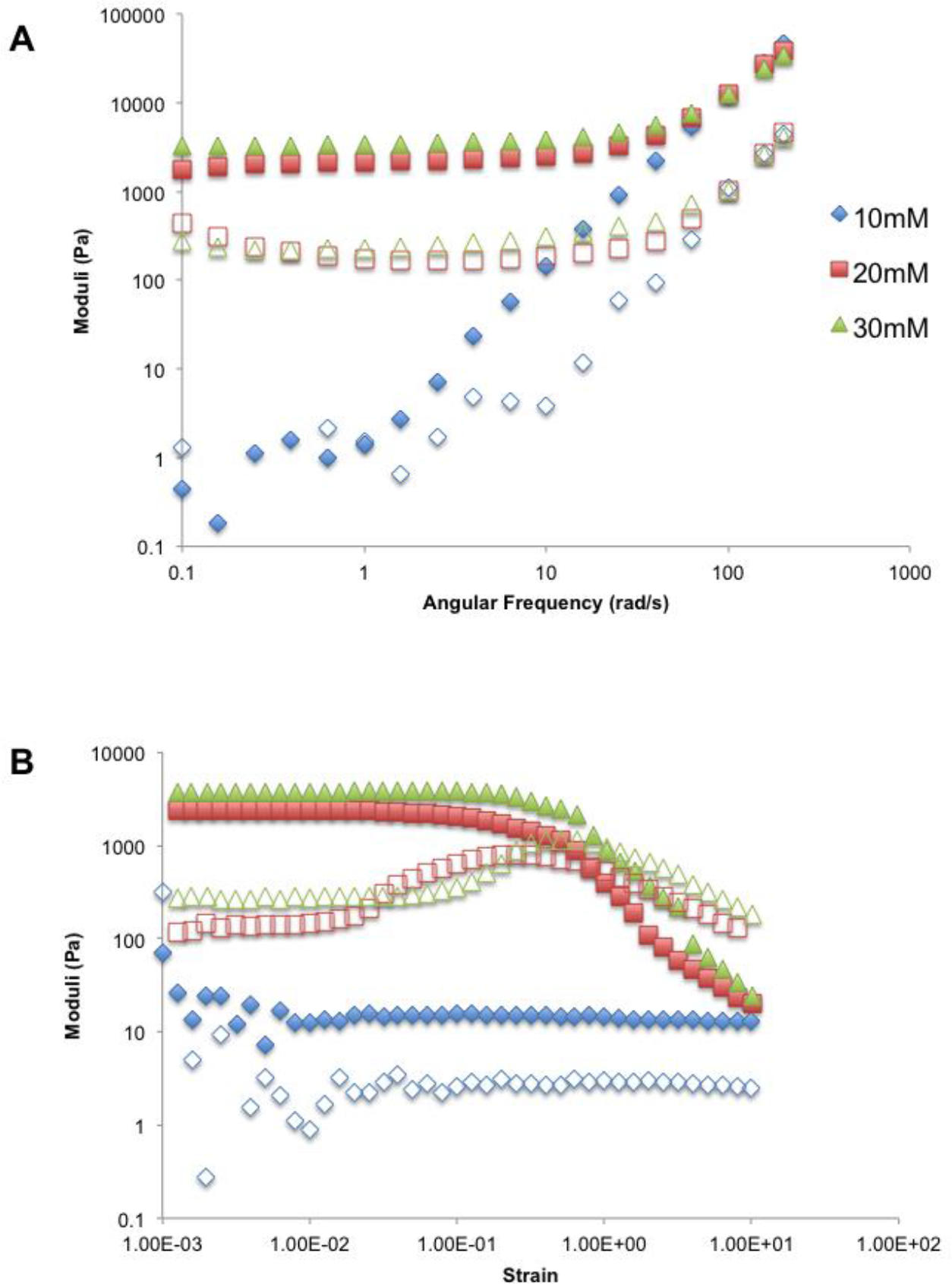
Representative frequency sweep and strain sweep curves of alginate hydrogels. A) Frequency and (B) strain sweeps for one set of alginate gels. Gels are 4% alginate with 10, 20, or 30mM CaCO_3_. Frequency sweeps were done at 1% strain and strain sweeps were done at 3.14 radians/s. The elastic moduli (G’) are shown with solid symbols and the viscous moduli (G”) are shown with hollow symbols of corresponding shape and color.

### Neutrophil Detachment and Engulfment of Pieces of Hydrogels

As an indicator of the success of neutrophils at breaking off and engulfing pieces of gel, we used phase contrast and epifluorescence microscopy to determine the percentage of neutrophils that, after one hour’s exposure to the gel, contained fluorescent beads that had originally been embedded within the hydrogel. For this, at least 10 fields of view under the microscope were chosen at random, and at least 100 neutrophils were counted, per sample. At least 3, and up to 5, replicate experiments were done, on different days, for each agarose or calcium concentration. The fluorescent beads were under 1μm in size and a much lower volume fraction of the gel than the polymers themselves so they would not be the predominate factor in determining mechanics of the gels. The hydrogels were in the bottom of 24-well plates and much larger than the neutrophils themselves, therefore, the relative shape compared to the neutrophil would be a flat surface representative of a much larger spherical target.

#### Agarose

The percentage of neutrophils containing beads engulfed from agarose gels decreases sharply upon changing agarose concentration from 0.3% to 0.5% (Figure 3). For 0.3% agarose gels, on average 28% of neutrophils had internalized fluorescent beads. For agarose gels at 0.5% agarose and above, on average less than 2% of neutrophils contain the fluorescent beads that are the marker for successful engulfment of gel fragments.

**Figure 3.**
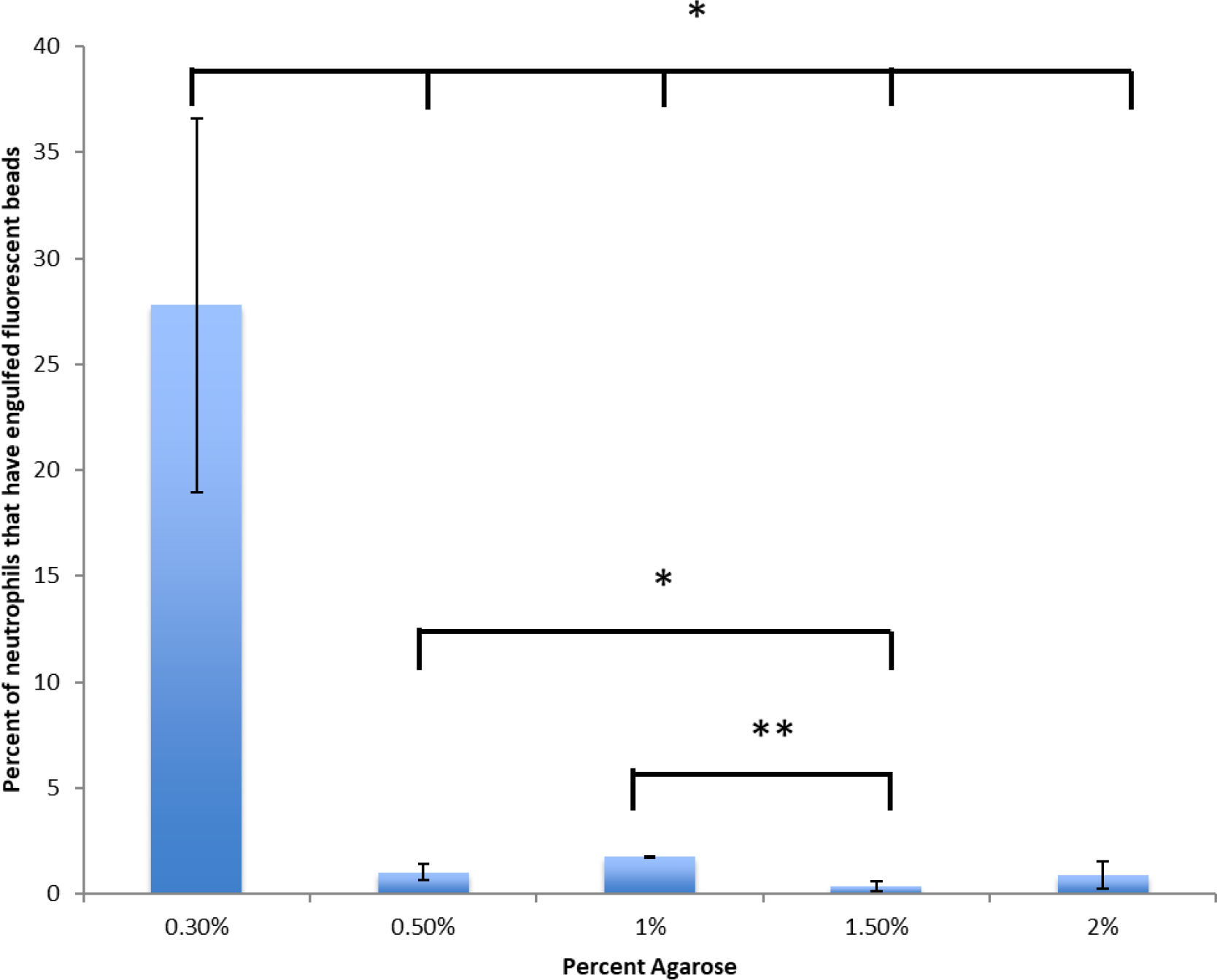
Neutrophils more successfully detach and engulf pieces of low-concentration agarose gels. At 0.3% agarose, an average of 28% of neutrophils had internalized beads. For all other agarose concentrations, fewer than 2% of neutrophils had internalized beads. Error bars are standard error of the mean. One * indicates a p-value between 0.05 and 0.005. Two ** indicates a p-value of less than 0.005, determined by a two-tailed Student T test.

#### Alginate

For alginate gels cross-linked with 10 mM calcium, on average, 77% of neutrophils engulfed beads initially contained in the gel. (Figure 4 and 5). For alginate gels cross-linked by 20mM calcium, on average 5% of neutrophils engulfed beads, and for alginate gels cross-linked by 30mM calcium, on average fewer than 2% of neutrophils engulfed beads.

**Figure 4.**
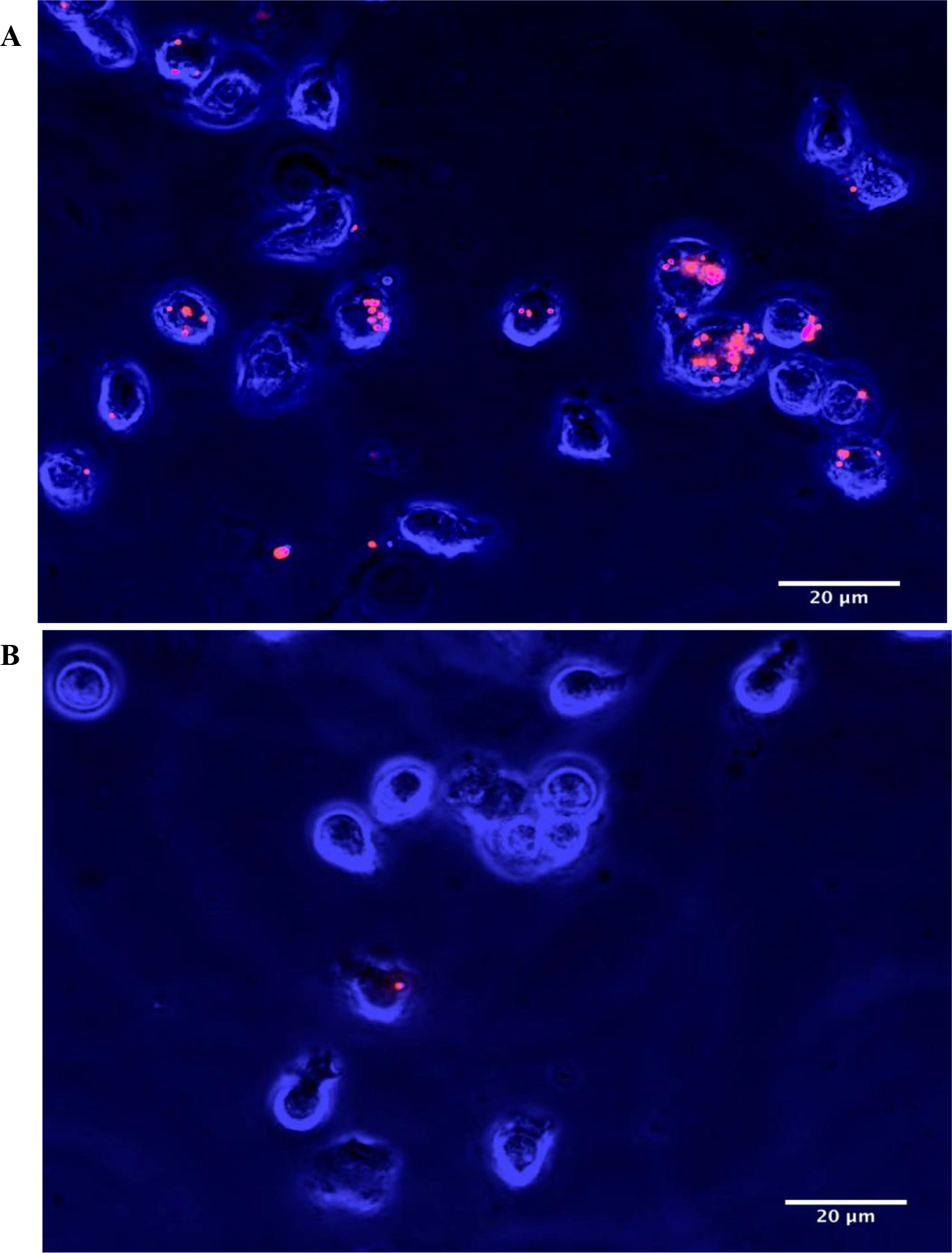
Neutrophils with internalized fluorescent beads from alginate gels. False-colored phase contrast images of neutrophils (blue) combined with fluorescence images of beads (red) after incubation with alginate gels cross-linked with 10mM calcium (A) and 30mM calcium (B).

**Figure 5.**
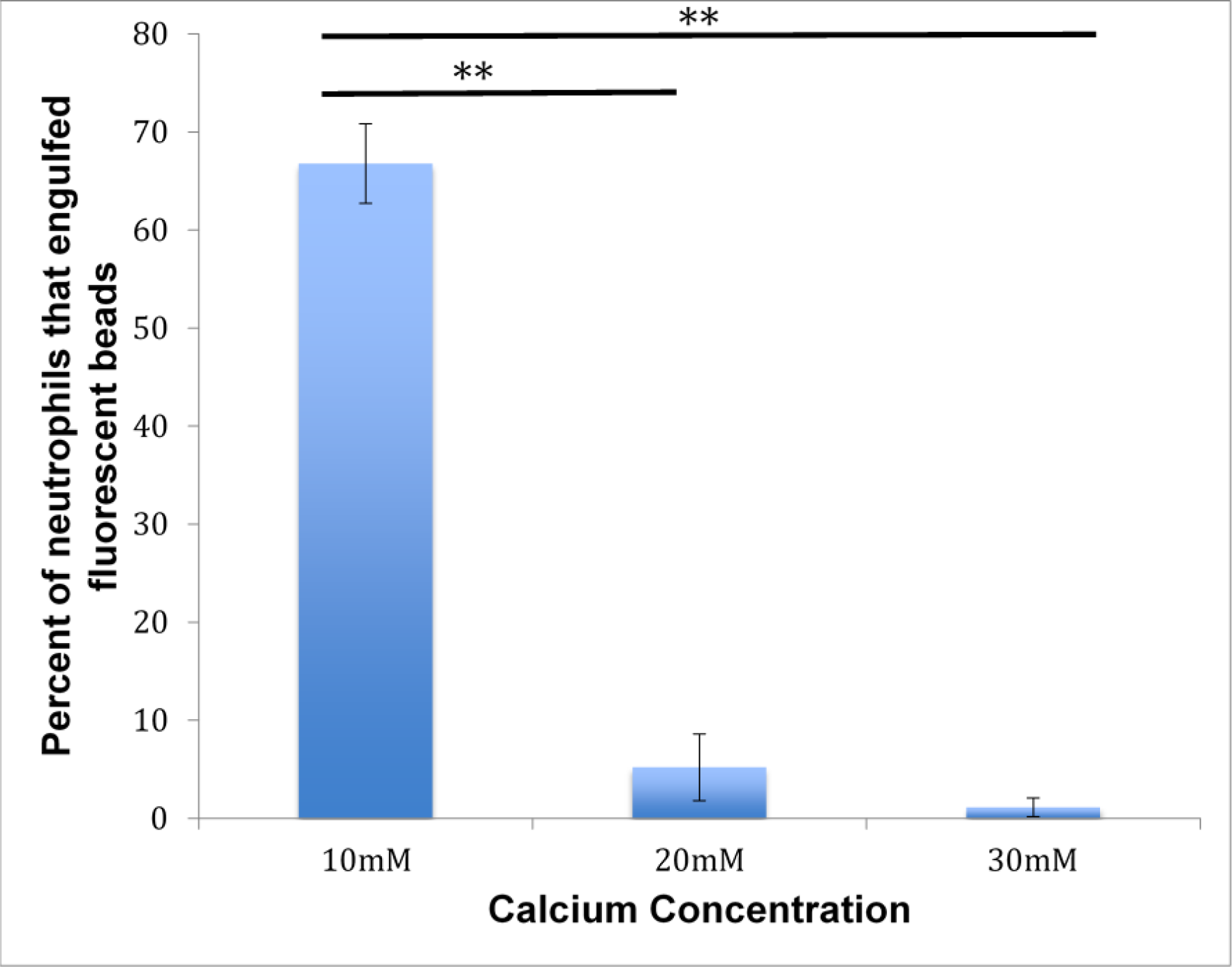
Neutrophils more successfully engulf alginate gels cross-linked with lower concentrations of calcium. As the concentration of calcium increases and therefore calcium cross-linking increases, the ability of neutrophils to successfully detatch and engulf pieces of the gels, and thereby and internalize beads, decreases. Error bars are standard error of the mean. One * indicates a p-value between 0.05 and 0.005. Two ** indicates a p-value of less than 0.005, determined by a two-tailed Student T test.

### The Success of Neutrophils at Engulfing Parts of Large Hydrogel Targets Depends Negatively on the Elastic Modulus of the Target

Using both alginate and agarose gels allows an initial disentangeling of the effects of elastic modulus from any effects arising from specific chemical composition. For both hydrogels, we find that the percent of neutrophils that successfully engulfed beads varies with the elastic modulus in such a way that data from the two types of gel lie very nearly along the same curve (Figure 6).

**Figure 6.**
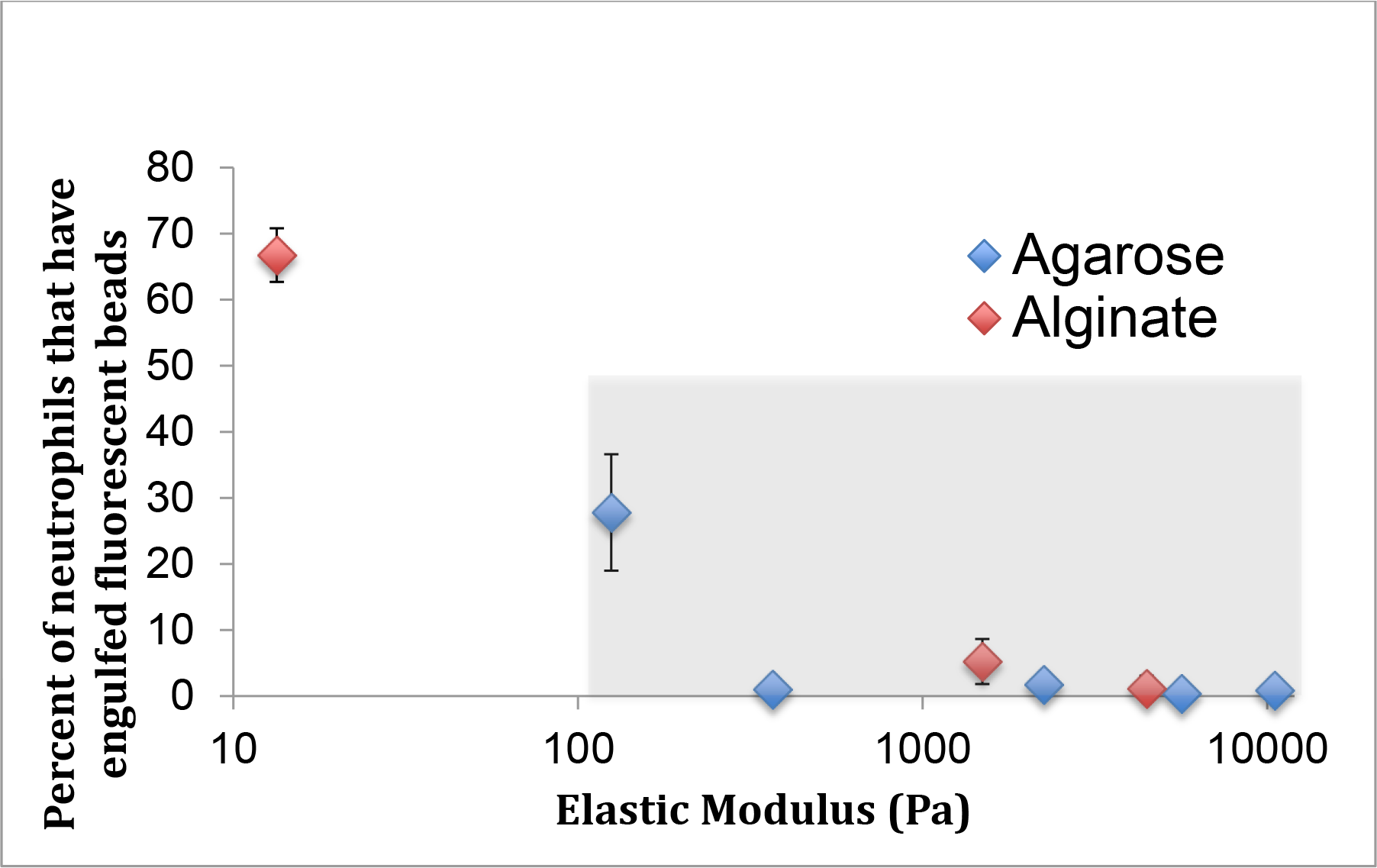
Percent engulfment lowers as elastic modulus of hydrogel increases. Elastic modulus includes moduli measured from both alginate and agarose, as indicated in the figure legend. Data points from alginate engulfment experiments are in red and agarose experiments are blue. There is a general trend of a decrease in the number of neutrophils that have internalized beads as elastic modulus increases, regardless of hydrogel material. X axis in a log scale to aid in visualization of the lower elastic moduli points. The grey shaded region indicates the range of elastic moduli that we measured earlier for biofilms re-grown from clinical isolates taken from patients with Cystic Fibrosis ^1^.

## DISCUSSION

### Summary of Results

We have developed a straightforward and inexpensive assay for measuring how the viscoelastic mechanics of a large target impact the ability of phagocytic cells to remove and engulf part of the garget. We have demonstrated the utility of this assay by measuring the impact of elastic modulus on engulfment by neutrophils. We find that increasing the elastic modulus of a large, viscoelastic target decreases the ability of neutrophils to successfully engulf the target. The dependence of engulfment success on elastic modulus is very similar for both alginate and agarose hydrogels, indicating that mechanics, not material chemistry, is likely the dominant factor governing the success of phagocytic engulfment in these cases. The range of elastic moduli that we re-created in these gels includes the range of elastic moduli that we previously measured for biofilms grown from clinical bacterial isolates, ~0.05 – 10 kPa ^1^. This range is indicated by the grey box in Figure 6. For gels with moduli spanning this range, we find that the success of neutrophils at detaching and engulfing parts of the gel falls from about 30% success to about 0% success. This suggests that the elastic modulus of biofilm infections might impact their resistance to the immune system, and that the evolutionary trend toward promoting biofilm toughness that we showed in our earlier work may reflect a selective advantage of higher elasticity for biofilms.

### Other Potential Influences on Neutrophil Engulfment

The studies in this paper focus exclusively on the elastic modulus of large, viscoelastic targets. However, the yield strain and yield stress, as well as other mechanical properties, likely influence the success of phagocytosis as well. Furthermore, the strength of the adhesion of the phagocytic cell to the target could also impact the success of phagocytosis. Future work could extend the method we present here to examine the impact of these other properties.

To engulf particles, neutrophils must bind to the target as it extends its cell membrane to engulf it. The more binding sites, the stronger a neutrophil can adhere and wrap itself around the target ^7, 33^. In our study, to make the abiotic gels recognizable to the neutrophils, bovine serum albumin (BSA) was added into the gel and was then incubated with an anti-BSA antibody before exposure to the neutrophils. Previous studies of the mechanical limitations of neutrophil phagocytosis have been measured only with antibody-mediated phagocytosis ^7, 33^. This method has been used to study neutrophil mechanics when the target is a stiff, polystyrene bead ^7, 33^. In our studies, we saw that the presence or absence of antibodies did not affect percent engulfment of alginate gels. BSA was still added to the gels, but as engulfment of the gels occurs without the antibodies present, phagocytosis of the abiotic alginate and agarose gels is not necessarily mediated through the Fc receptors.

### 4.3.3 Health Implications

The increased resistance to phagocytosis conferred by increased elastic modulus of biofilms could result in worse outcomes for infected patients if it causes frustrated phagocytosis, in which neutrophils release reactive oxygen species that damage host tissue as well as bacteria ^34, 35^. Indeed, damage from the patient’s own inflammatory response and associated release of reactive oxygen species is the primary cause of lung failure in cystic fibrosis patients ^36, 37^. Mechanical resistance to phagocytosis could also allow more time for bacterial virulence factors, such as pyocyanin and rhamnolipids produced by *Pseudomonas aeruginosa*, to be produced and to damage neutrophils ^38–40^.

### Conclusions and Future Work

We have developed a method for determining how the viscoelastic mechanics of a large target impacts the phagocytic success of an attacking cell. Using this method, we have shown that sufficiently-high elastic moduli can limit or prohibit the success of phagocytosis. Future work could extend this method to account for the effect of other mechanical properties of the target, such as yielding, and of the mechanical forces binding the phagocytic cell to the target. The use of gels with specifically-tunable yield strains and surface chemistries, as well as elastic moduli, would be desirable for this. Our study here measures phagocytic success only after a fixed time, but our method is readily extensible to time-varying studies to determine how mechanics impacts the timescale for phagocytic success. It is known that the shape and initial angle of contact between a macrophage and its target determines whether or not phagocytosis is successful ^9^, but this has yet to be confirmed in neutrophils.

Using the method we present here, and extending these studies as outlined in the preceeding paragraph, we expect to be able to determine what mechanical properties render a biofilm most susceptible to phagocytic clearance. We expect that this will guide us and other researchers as we work to develop new, non-antibiotic approaches to biofilm treatment, which we hope will circumvent biofilm’s innate, phenotypic resistance to antibiotics, reduce damage to patients’ health by the harmful side effects of antibiotics, and slow the evolutionary development of antibiotic resistance.

## Methods

### Hydrogel preparation

Abiotic hydrogels were produced using alginic acid sodium salt powder or low-gelling temperature agarose (both purchased from Sigma-Aldrich).

Alginate gels were made by dissolving 4% sodium alginate in distilled water and subsequently adding 5% calcium carbonate (CaCO_3_) and 5% D-(+)-Gluconic acid δ-lactone (GDL). Alginate gels were made using 10, 20, and 30mM CaCO_3_; Ca^2+^ acts as a crosslinker.

By increasing the length of the G-block of alginate, mechanical properties can be enhanced ^41^. The ratio of M:G residues varies across species that produce alginate therefore mechanics will vary for alginate gels depending on the source of alginate. We used sodium alginate isolated from brown algae and while it is a cheaper, more accessible source of alginate, its G-block composition is more variable than others. Alginate gels consistently were made of 4% alginate and 5% calcium carbonate solution, with the concentration of CaCO_3_ in the solution varying to change the overall calcium concentration. 5% GDL was added to cause internal gelation by slowly lowering the pH of the gel causing calcium ions to be released from CaCO_3_ into solution ^42^.

Agarose gels were made using low-gelling temperature agarose at 0.3%, 0.5%, 1%, 1.5%, and 2%.

Both alginate and agarose gels contained 10mg/ml bovine serum albumin (BSA) (HycClone brand, purchased from GE Lifesciences) for immune activation and a 1:100 dilution of fluorescent beads (0.955μm polystyrene Dragon Green beads, purchased from Bangs Laboratories) for microscopic visualization. Hydrogels used for microscopy experiments were stored overnight at 4°C in 500μl aliquots in 24-well flat-bottom plates. Hydrogels used for rheological measurements were poured into small petri dishes at 1000-2000μm deep and stored overnight at 4°C.

### Rheology

We measured hydrogel bulk mechanics using oscillatory bulk rheology done by a stress-controlled AR 2000ex rheometer from TA Instruments. Hydrogel samples were taken out of 4°C storage on the morning of the experiment and placed at 37°C for one hour. Then, hydrogel sections were placed on the rheometer and cut down to the size of the 8mm parallel plate head. The gap height varied between 1 and 1.5mm depending on the specific gel sample. Oscillatory frequency sweeps from 0.1 to 600 rad/s at 1 % strain and strain sweeps from 0.1 to 200 % at 3.14 rad/s were performed on each sample on each day of measurement. Elastic modulus was taken as the midpoint of the G’ plateau region for strain sweeps.

### Neutrophil Isolation

Human neutrophils were isolated from two adult volunteer blood donors following a protocol published by others ^43^. This and the rest of the work on neutrophils was approved by the Institutional Review Board (IRB) at the University of Texas at Austin as Protocol Number 2015-05-0036.

Unless otherwise specified, materials were purchased from Sigma-Aldrich. In brief, blood was collected in heparin-coated tubes, mixed with a 3% dextran and 0.9% sodium solution, and red blood cells fell out of solution. The remaining supernatant was centrifuged for 10 minutes at 500g and the resulting pellet was re-suspended in 10ml of Hanks Buffered Salt Solution (HBSS) without calcium or magnesium. Cells were separated using a Ficoll-Paque density gradient solution (purchased from GE Healthcare), spun at 400g for 40 minutes. This resulting pellet was re-suspended in distilled water for 30 seconds to lyse any remaining red blood cells. Then, the isotonicity of the solution was restored using 1.8% NaCl. Cells were centrifuged for 5 minutes at 500g and the final neutrophil pellet was re-suspended in 1ml HBSS with calcium and magnesium and 20% human serum.

### Neutrophil Engulfment Assay and Microscopy

On the day of phagocytic engulfment experiments, neutrophils were isolated as described above. While neutrophils were being isolated, 100μl of rabbit anti-BSA antibody diluted 1:1000 in Dulbeccos’ phosphate-buffered solution (DPBS) (Invitrogen brand, purchased from Thermo-Fisher Scientific) was added to each hydrogel and incubated at 4°C for 30 minutes. Hydrogels were then washed three times with DPBS. After neutrophil isolation was complete, 200μl of cell suspension in HBSS and serum was added to each well. The hydrogel plate with neutrophils was then incubated at 37°C for one hour; 37°C is human body temperature. After one hour, the solution containing cells was collected from off the top of the hydrogel. In some cases, to increase the number density of cells in the microscope field of view, collected neutrophils were concentrated and re-suspended in 50μl of HBSS. The collected cells were put on a microscope slide with a coverslip that had been coated in0.1% poly-l-lysine (Sigma-Aldrich). Cells were imaged using phase contrast microscopy and beads were imaged using epifluorescence microscopy with a filter for green fluorescent protein. At least 100 neutrophils from every sample, from fields of view chosen at random were counted as either containing or not containing fluorescent beads, which served as indicators of gel engulfment.

### Growth of Bacterial Aggregates

Overnight shaken cultures of *P. aeruginosa* naturally contain both multicellular aggregates and single cells ^44^, and aggregates analyzed in the experiments formed spontaneously during overnight growth. To promote the formation of more and larger aggregates, we used a bacterial strain that over-expresses the extracellular polysaccharide Psl, ΔwspF Δpel ^45^. We grew aggregates using the same process we have described previously ^46, 47^, as follows:

Frozen bacterial stock was streaked onto a Lysogeny Broth (LB) agar petri plate and incubated at 37 °C overnight. From this plate, one colony was dispersed into 4mL of LB liquid growth medium and left to overgrow for approximately 20 hours at 37 °C on a rotating shaker. Cultures were overgrown to stationary phase to increase the number of large multicellular aggregates present.

### Microscopy of Neutrophils and Bacterial Aggregates

Freshly isolated human neutrophils at a concentration of ~10^6^ cells/mL were mixed with bacterial culture (diluted to OD_600_ = 0.2 measured using a Genesys spectrophotometer) and 20% human serum. The mixture of bacteria, which included both single cells and aggregates, and neutrophils, was then placed on a coverslip chamber to be imaged. Timelapse videos were obtained using an inverted phase contrast microscope with 60X objective (both from Olympus), QImaging Exi Blue CCD camera, and QCapture Pro 6 software. As described in our previous work, the stage region of this microscope is enclosed in an incubator chamber ^45, 48, 49^. During image acquisition the microscope stage incubator was set to 37 °C, which is human body temperature. Images were analyzed using Fiji, a software distribution of ImageJ.

### Author Contributions

M. D.-F. performed experiments, analyzed data, and wrote the paper. L. B. performed experiments, analyzed data, and wrote the paper. K. K. performed experiments. V. D. G. designed research and wrote the paper.

## Acknowledgements

We thank Prof. Nathaniel A. Lynd (Department of Chemical Engineering, The University of Texas at Austin) for the use of his rheometer and for helpful discussions. We thank Prof. Laura Suggs (Department of Biomedical Engineering, The University of Texas at Austin) for helpful discussions and assistance with the alginate gel. Neutrophil work was approved by the Institutional Review Board (IRB) at the University of Texas at Austin as Protocol Number 2015-05-0036. This work was supported by grants from the Cystic Fibrosis Foundation (Gordon 201602808-001), the National Institutes of Health (1R01AI121500-01A1, NIAID), and the National Science Foundation (727544, BMMB, CMMI), all to Vernita Gordon.

## Supplementary Movie Caption

These phase contrast micrograph movies show neutrophils interacting with and attacking biofilm-like aggregates of bacteria at 8 times real speed. Each has a 30 μm scalebar. Aggregates are seen as the bacteria-dense regions that exclude biofilms. Aggregated bacteria are bound together in a biofilm-like matrix of polymer and protein which is not visible under phase contrast microscopy.

